# Posterior inferotemporal cortex cells use multiple visual pathways to complement fine and coarse discriminations

**DOI:** 10.1101/048561

**Authors:** C. R. Ponce, S. G. Lomber, M. S. Livingstone

## Abstract

In the macaque monkey brain, posterior inferior temporal cortex (PIT) cells are responsible for visual object recognition. They receive concurrent inputs from visual areas V4, V3 and V2. We asked how these different anatomical pathways contribute to PIT response properties by deactivating them while monitoring PIT activity. Using cortical cooling of areas V2/V3 or V4 and a hierarchical model of visual recognition, we conclude that these distinct pathways do not transmit different classes of visual features, but serve instead to maintain a balance of local-and global-feature selectivity in IT.

## INTRODUCTION

Posterior IT (PIT) neurons are the penultimate stage of the ventral visual processing stream, comprising cortical areas V1→V2→V4→PIT→anterior IT (AIT). In addition to this main pathway, PIT also receives direct feedforward projections from V3 and V2^1^, and V4 receives direct inputs from V1. These short projections (V1→V4→PIT and V1→V2→PIT) have been called bypass pathways^2^, and they represent a significant fraction of the inputs to PIT: 14% of all neurons in the brain projecting to PIT are located in areas V2|3 (for context, 26% of inputs to PIT arrive from V4; only 1% of inputs to V1 come from the LGN)^3^,^4^. The remaining projections arise from AIT and the dorsal pathway. The goal of this study is to define the role of these different input pathways in PIT selectivity.

We recorded from an unbiased sample of PIT neurons while deactivating, by cooling, areas V2−V3 (together) and V4 (**Fig. 1a**). We measured PIT firing rates before and during cooling of V4 or V2|3, and quantified changes in the representational capacity of PIT. Using firing rate statistics and linear classifiers (i.e. support vector machines), we found that while V4-dependent inputs were more important for preserving the representation of the identity of images in PIT, the different concurrent pathways did not transmit different types of visual features (such as different proportions of curvature or spatial frequency). We modeled the contributions of short-and long-pathways using the standard model of visual recognition^5^, and observed that short pathways were well-positioned for fine feature discriminations. We confirmed that fine-feature discrimination was relatively better preserved during cooling compared to coarse-feature discrimination. We simulated the effects of cooling on decoding accuracy using various random cooling effects models, and found that while these random models predicted an overall loss of accuracy, they did not predict the preservation of decoding accuracy for fine discriminations. Thus we conclude that short pathways are helpful in fine discrimination, because their receptive field weights match simpler image elements. By introducing units with simpler preferences into PIT, the short pathways create a diversity of feature preferences available for downstream perceptual operations.

**Figure 1.**
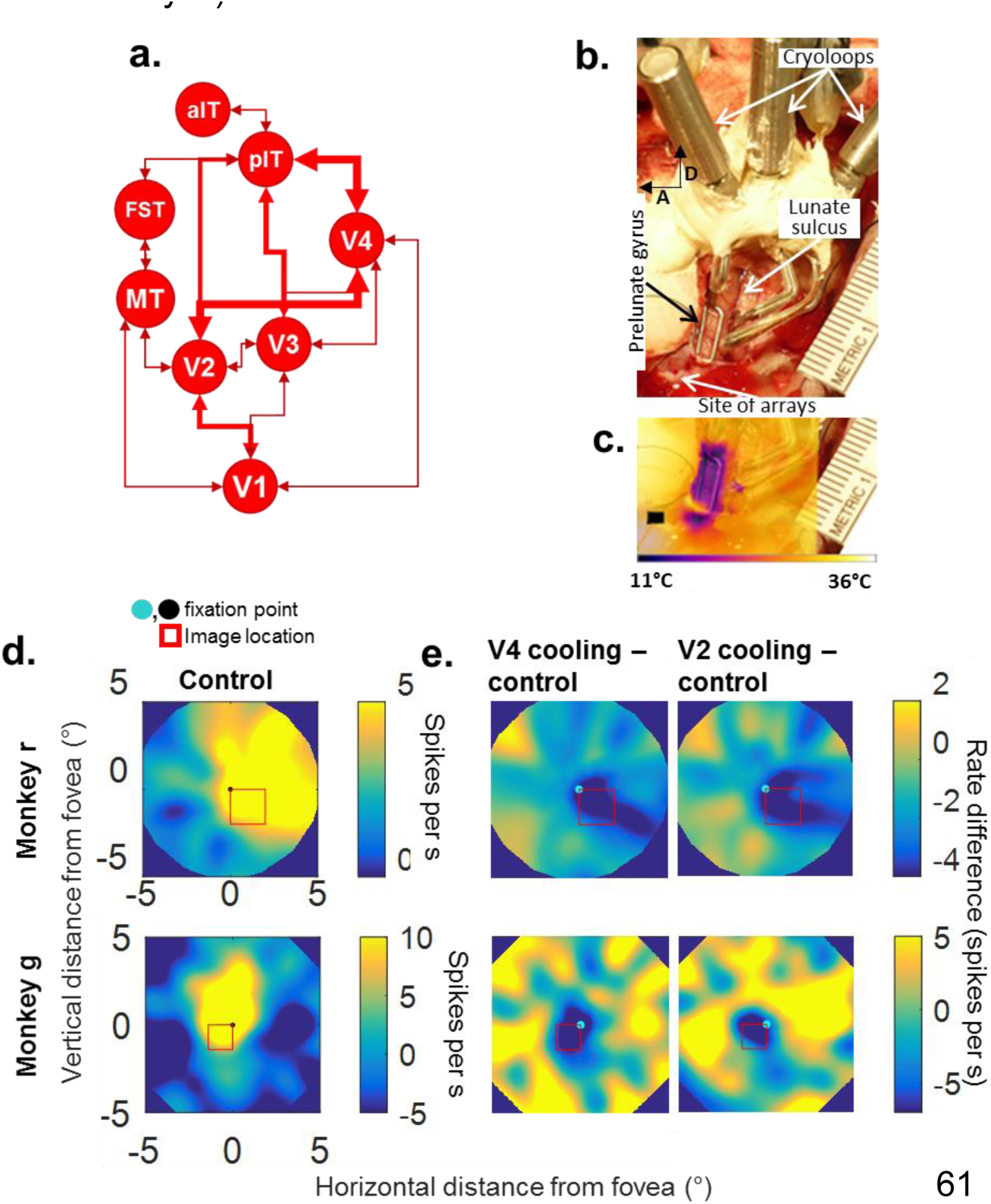
Cooling affected portions of the aggregate PIT response fields. **a.** Input network to posterior IT (PIT). **b.** Intraoperative picture, monkey R left hemisphere. Posterior craniotomy showing the location of the lunate sulcus, prelunate gyrus and V2/V4 cryoloops. The arrays were later implanted where shown by the label. A: anterior, D: dorsal. **c.** Composite image showing the superimposed thermal and visible light images, while the prelunate gyrus loop is active. The black square shows the approximate location of the arrays. **d.** Average firing rate for all units, evoked by flashing an image in a 8 × 8° grid, during the control (warm) condition. **e.** Difference in activity during cooling of area V4 (left column) or V2|3 (right column) – the control map was subtracted from the cooling map. Dark regions show reductions in firing rate.

## RESULTS

### Cooling affected portions of PIT response fields

We implanted floating microelectrode arrays in the PIT of two adult male monkeys (2 arrays in monkey R and 3 arrays in monkey G), along with cryoloops on dorsal V2, V3 and V4^6,7^. The arrays were placed anteriorly to the inferior occipital sulcus, the cryoloops were placed within the lunate sulcus and over the predorsal gyrus (**Fig. 1b**). We activated the cryoloops intraoperatively, using thermal imaging to plot the extent of cooling and found that the lower thermal region was limited to 1-1.5 mm around and within the cryoloop (**Fig. 1c**). The flat area of cortex directly cooled by the cryoloops was around 13 × 5 mm^2^. The electrode arrays were at least 5 mm anterior to the prelunate cryoloop, and anterior to the inferior occipital sulcus. Three weeks after each surgery, we measured the retinotopic response fields of the arrays and the retinotopic size of the cooling scotomas: the animals maintained their gaze on a 0.4°-diameter central black circle, while a 2°-diameter image was flashed randomly within a 16 × 16° radial grid. We collected spike data before, during and after activation of the V2/V3 or V4 cryoloops, counterbalancing the order of the V4 vs. V2|3 deactivations. The mean PIT population response fields were centered on the perifoveal upper contralateral hemifield (**Fig. 1d**). In contrast, the V4 and V2|3 scotomas were centered towards the perifoveal lower hemifield, as predicted by the dorsal retinotopic representations of V2, V3 and V4. In subsequent experiments, stimuli were sized to fit within the overlapping region of both scotomas (**Fig 1e**) (1.4°-wide images for monkey G, 2.0°-wide images for monkey R).

### Cooling reduced firing rates in PIT units and decoding accuracy by linear classifiers

In all the following experiments, we showed the fixating animals 293 images belonging to 15 different categories (angles, animals, artificial objects, curves, faces, radial and linear gabors, joint angles, plants, places, noise textures and tristars; the entire image set is shown in **Supplementary Fig. 1**). When we cooled either set of coils, PIT multiunits reduced their visually evoked spike rates (**Fig 2a**). During control conditions, PIT multiunits showed a median visual response of 18±1 (monkey R) and 22±2 (Monkey G) spikes/s (range of −3 to 120 spikes per second relative to baseline for monkey R; −12 to 106 spikes per s for monkey G). When the V2/V3 loops were cooled, the overall average rate was reduced to 13±1 and 14±1 spikes/s (monkeys R, G). When the V4 cryoloops were cooled, the overall rate was reduced to 12±1 and 15±1 spikes/s (monkeys R, G; probability that all means come from same distribution *P* = 6×10^−4^ and 1×10^−4^, one-way ANOVA, N_sites_ = 300 and 256, F(2, 897) = 8.1, F(2,765) = 8.9). In one animal, we cooled both V4 and V2|3 loops concurrently, measuring a similar reduction in firing rate (38%); cooling both sets of loops did not silence PIT activity. Another measure of input strength is response latency, and here we similarly observed little difference between V2/V3 cooling and V4 cooling (see **Supplementary Section 1**).

**Figure 2.**
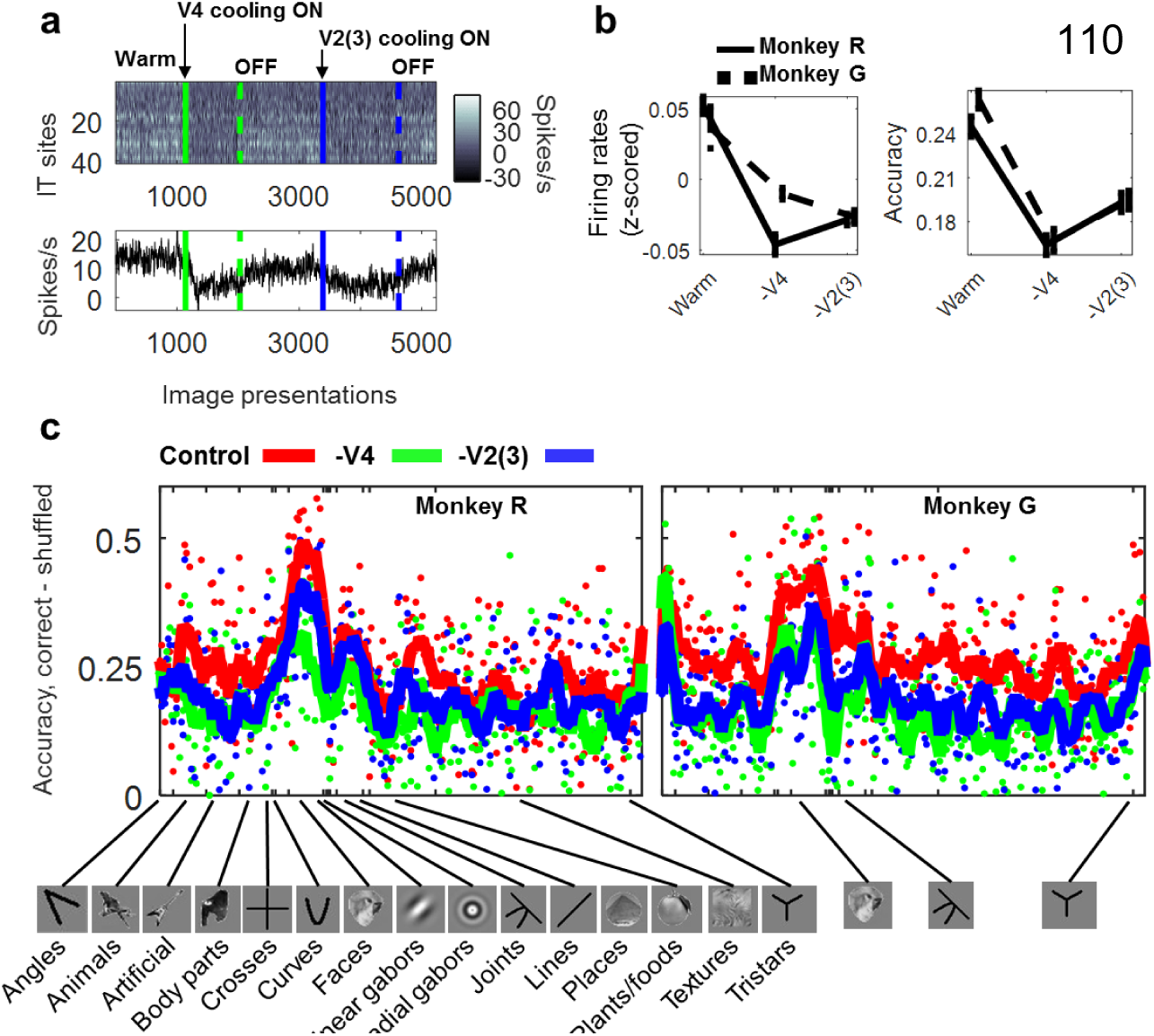
Effects of cooling on firing rate and classification accuracy. **a.** (top) Data from one cooling session (monkey R, day 1). The top figure shows the evoked spike rates from 50 PIT sites (rows) recorded concurrently before, during and after cooling of V4 and V2|3. Each column represents one image presentation. Solid lines mark the onset of each cooling condition, broken lines show the onset of the rewarming periods. (bottom) Mean firing rate across each temperature condition. **b. (top)** Average firing rate activity (z-scored) for all channels during each temperature condition. **(bottom)** Average classification accuracy for all images during each temperature condition. **c.** Mean cross-validated accuracy for each image before and during cooling (red = control, green = V4 cooling, blue = V2|3 cooling). The x-axis shows all 293 images listed alphabetically by their category.

Next we used pattern analysis to quantify the encoding capacity of PIT during V4 or V2|3 cooling. We trained statistical classifiers (support vector machines with a linear kernel, or SVMs) using data from each experimental condition (before cooling, during V4 or V2|3 cooling). SVMs were used in an all-vs.-all approach, so chance performance was 50% per comparison. We had few trials per image during each cooling condition (4-6) and so we used leave-one-out cross-validation for each paired comparison. To further guard against unreliable values due to the small sample number, we also trained SVMs using the same data but shuffling the image labels. Thus, accuracy was defined as the mean of all cross-validation cycles using the correct labels minus the mean of cross-validation cycles using the shuffled labels, so a range of 0-0.5 is equivalent to 50-100% accuracy.

First, this decoding analysis showed that faces elicited the highest classification accuracy in both animals, which was notable because we did not pre-select the array implantation sites by proximity to fMRI-defined face patches. SVMs showed a median accuracy value of 0.25±0.01 and 0.26±0.01 above baseline (monkeys R, G, standard error of the median). During V4 deactivation, median accuracy dropped to 0.16±0.01 and 0.17±0.01; during V2|3 deactivation, median accuracy dropped to 0.19±0.01 and 0.20±0.01. These median accuracy values were statistically different at the group level (monkeys R, G: *P* = 5×10^−12^, 3×10^−23^, one-way Kruskal-Wallis, N_*images*_ = 347, 293, χ^2^ (2,1038)=52, χ^2^(2,876)=104). The differences in median values between V4 and V2|3 cooling were statistically reliable in both animals (monkeys R and G: *P* = 2×10^−5^, 4×10^−3^, two-tailed Wilcoxon sign rank test, N = 347, 293 accuracy values, Z-stats: −4.3, −2.9, **Fig. 2b**, bottom, and **Fig. 2c**). We also performed an SVM category classification analysis, grouping responses by category, not as individual images. We also found that deactivating V4-inputs reduced category classification accuracy more than did deactivating V2|3 (**Supplementary Section 2, Fig. S2 and S3**).

In summary, PIT multiunits showed similar mean firing rate reductions during V2|3 or V4 deactivation, but in both monkeys, SVM accuracy was reduced more by V4 deactivation. This suggests that V4 direct inputs are more important for image identification and categorization, and that this cannot be explained by a simple reduction in mean spike rate. We further explored the reasons why decoding accuracy shows a quantitatively stronger role for V4 using a multidimensional projection analysis, described in **Supplementary Section 3.**

### Cooling effects on median accuracy were simulated by random processes

To gain a better understanding of the drop in decoding performance during input cooling, we asked how much of the accuracy impairment could be attributed to non-specific fractional reductions in firing rate. We modeled three mechanisms of cooling rate reductions: 1) each IT multiunit underwent a given fractional reduction across all of its responses (a *site-by-site* cooling mechanism); 2) all multiunits underwent the same fractional reduction, but the reduction could vary over time (a *temporal* mechanism); finally, 3) each multiunit underwent a mix of *site-by-site* and *temporally dependent* reductions. We first examined the effects of each cooling mechanism using a model. In neural activity space, where images are represented as coordinate positions in a multi-dimensional space (**Fig. 3a**), these cooling effects would induce different geometric transformations and thus affect decoding accuracy differently. Consider the first mechanism of site-by-site cooling: this has no effect on classification accuracy, because each site’s responses are normalized before classification; it makes no difference if the lengths of the population response vectors change, as long as they keep the same direction (**Fig. 3b-c**, ***i***). This is a plausible compensation because normalization is a common mechanism in cortical computations. The second mechanism, temporally dependent reductions, where cooling imposes a different fractional value on the whole population vector at different times, pulls and stretches the pre-cooling response vector groups towards a minimum (**Fig. 3b-c**, ***ii***). The third mechanism, where individual multi-unit sites undergo different fractional reductions over time, is interesting because it changes the direction of the vectors, directly affecting the image representations in activity space (**Fig. 3b-c**, ***iii***).

**Figure 3.**
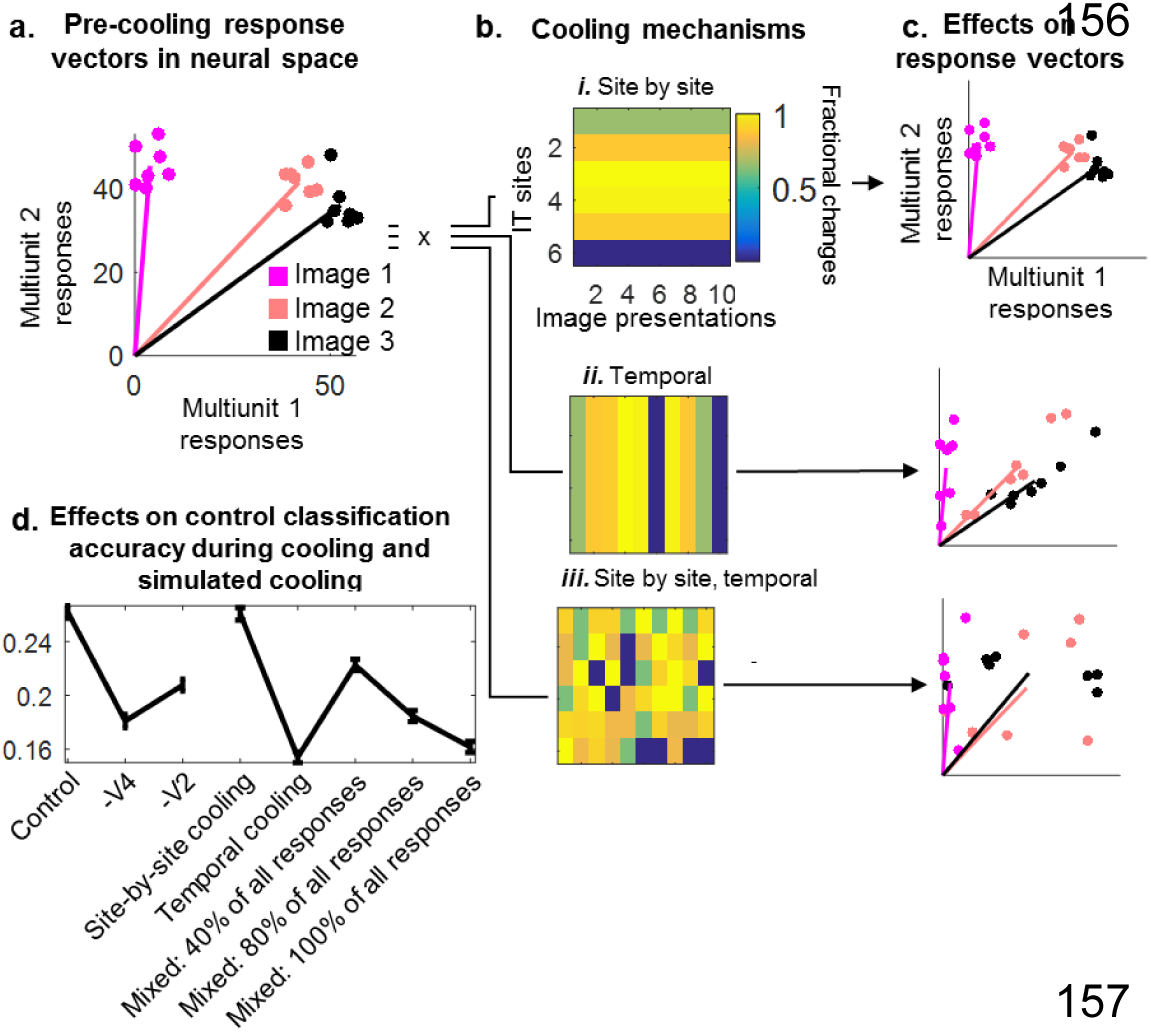
Cooling simulations. **a.** Hypothetical responses of two units (axes) to three different images (colors), each presented seven times each (points). **b.** Different mechanisms of cooling. Cooling may impose a fixed fractional change in a channel-by-channel basis, a temporal basis, or a combination of the two. **c.** Responses in (A) transformed by each “cooling” reduction rate mechanism. **d.** Mean accuracy values (baseline-subtracted, ± standard error) measured before cooling (“control,” “−V4,” “−V2|3”, both monkeys combined), and after different cooling simulations (“site-by-site”, “temporal”, and “mixed”). For the “mixed” simulations, each percentage shows the fraction of responses that were randomly multiplied by the gain values.

We applied each mechanism to our control (warm) data to simulate the cooling drop in decoding performance. First we measured the distribution of fractional cooling changes during cooling. In monkey R, the median fractional reduction during V4 cooling was 0.68 (25th, 75th quantiles: 0.51-0.80), and during V2|3 cooling, 0.73 (25th, 75th quantiles 0.59-0.84). For monkey G, the median fractional reduction during V4 cooling was 0.67 (25th, 75th quantiles 0.54-0.82), and during V2|3 cooling 0.62 (25th, 75th quantiles 0.48-0.74). Note that some of these fractional changes included increased firing rates during cooling, but this was expected from weakly firing multiunits. Next, we sampled these fractional distributions (with replacement) and multiplied each sampled fraction times the control (warm) response counts. These multiplications were done using either the site-by-site cooling mechanism, the temporally dependent cooling mechanism, or the mixed mechanism. As a form of cross-validation, the fractional gain distributions came from different days from the control data. We created 20 “cooling” populations, based on V4 and V2|3 cooling, per monkey.

Before cooling, the mean decoding accuracy for both animals was 26±1% over baseline, 16±1% during V4 cooling and 19±1% during V2|3 cooling. We found that the site-by-site spike mechanism did not reduce decoding accuracy (its value remained 26±1%, same as control). The temporally dependent mechanism reduced decoding accuracy to 15±1% over baseline. The mixed mechanism lowered accuracy as a function of the number of responses that were randomly affected: if we multiplied 100% of all responses by the random fractions, decoding accuracy was reduced to 16±1%. To match the experimental reductions in accuracy, we had to affect between 40-80% of all responses, which resulted in 18-22% decoding accuracy, **Fig. 3d**). In summary, we could not account for the observed reduction in decoding accuracy by a uniform fractional in spikes within each multiunit site, but a temporally varying reduction or a mix of site-by-site and temporally varying reductions could account for the mean decoding accuracy loss during cooling.

### Cooling reduced selectivity of individual PIT multiunits

Cooling inputs to PIT reduced classification accuracy at the population level. To examine accuracy at the level of individual sites, we measured selectivity using an F-test. Let us say that PIT units were selective to specific images if the mean variance of their spike counts to different images was greater than the mean variance of their spike counts to each image (F-statistic); the value 1 suggests no selectivity; the greater the value, the more selective. We called the F-test statistic per channel before cooling *F*_*control*_, during V4 cooling *F*_−*V*4_ and during V2|3 cooling, *F*_−*V*2|3_. If the distributions of *F*_−*V*4_ and *F*_−*V*2|3_ are closer to the non-selectivity value of 1 compared to the *Fcontrol* distribution, this would suggest that PIT sites became less selective during cooling.

Each PIT site gave one *F*_*control*_, one *F*_−*V*4_ and one *F*_−*V*2|3_ value. We plotted each control *F* value against its counterparts and found that cooling F-stats were mostly lower than the warm distribution (**Fig. 4c**, Monkey R, median *F* ± standard error, Warm: 1.13±0.03, −V4: 1.10±0.01, −V2|3: 1.14±0.02; Monkey G, Warm: 1.21±0.05, −V4: 1.13±0.02, −V2|3: 1.14±0.02). However, there was no statistical difference among these values (P = 0.20, 0.20, Kruskal-Wallis, N = 300, 256 sites, χ^2^(2,897)=3, χ^2^(2,765)=3). The reason there was no statistical difference is that many of the PIT sites were not that selective to start with, having pre-cooling *F* stats already bottomed out at 1. Still, units with higher pre-cooling F values showed greater changes during cooling. To quantify this observation, we asked if the slope describing the relationship between the pre-cooling and cooling *F* values was statistically different from unity. We used a bootstrap approach. For 1,000 iterations, we re-sampled sites with replacement and used their *F*_*control*_, *F*_−*V*4_ and *F*_−*V*2|3_ values to fit linear regression lines between control and −V4 values, and between control and −V2|3 values. This analysis resulted in 1,000 slopes describing the *F*_*control*_ and *F*_−*V*4_ relationship, and another 1,000 slopes describing the *F*_*control*_ and *F*_−*V*2|3_ relationship. None of these slope values overlapped the line of unity (Monkey R, mean slope ± SEM, control vs. −V4: 0.64±0.03, control vs. −V2|3: 0.82±0.04; Monkey G, control vs. −V4: 0.53±0.04, control vs. −V2|3: 0.59±0.03). We also noticed that the mean *F*_*control*_ / *F*_*cooling*_ slope was shallower during V4 cooling than V2|3 cooling in both animals. This implied that PIT multiunits became less selective during V4 cooling than during V2|3 cooling. This difference in slope was significant using a randomization test, which shuffled *F*_*cooling*_ values from the V4 and V2|3 conditions and asked if this mixed distribution could produce the observed slope (one-tailed randomization test, *P* = 0.001 and 0.02; see Methods for details).

**Figure 4.**
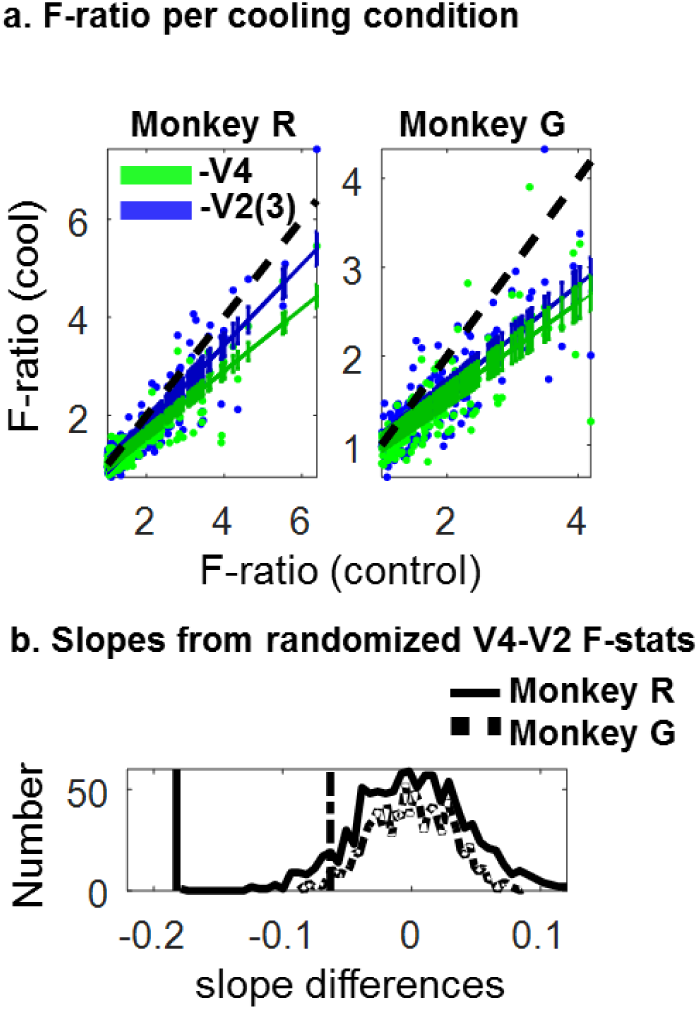
Selectivity of individual PIT sites before and during cooling. **(a)** Scatterplots of F statistics (warm vs. −V4: green, warm vs. V2|3, blue). Each point shows the paired F-ratios for a given channel, measured before and during cooling. The solid colored lines show the mean slope describing the control and cooling F-ratio distributions. The error bars around each slope show the standard error. The broken black line shows unity. **(b)** Differences in slopes expected given a mixed temperature distribution (solid curve = distribution from monkey R data, broken line = monkey G). The vertical lines show the experimental difference.

We further asked whether there was any relationship between the retinotopic location of a PIT response field relative to the cooling scotomas, and its subsequent selectivity (F-statistic) change. The images were presented at the intersection of the V4 and V2|3 scotomas. Therefore, some individual multiunit PIT response fields (RFs) would cover more of the stimulus than others. For each PIT site, we measured the fraction of its RF that overlapped the stimulus/scotoma, and correlated this value against the subsequent change in selectivity (F-statistic). The RF overlap measure was computed using data from different recording days. There was a small but statistically reliable correlation of RF overlap with selectivity change (selectivity change was defined as *F*_*control*_ - *F*_*cooling*_): during V4 cooling, the Pearson correlation coefficient was 0.19 and 0.32 (monkeys R and G: *P* = 1.2×10^−3^ and 1.3×10^−7^, Student’s t-test, *N* = 300, 256 sites). During V2|3 cooling, the correlation coefficient was 0.11 and 0.26 (*P* = 0.06 and 2.3×10^−5^). Note that the stimuli were placed in the same overlapping region between both −V4 and −V2|3 scotomas, so the lower correlation values for V2|3 cooling are not due to differences in scotoma overlap; rather it is because the selectivity change is less pronounced for V2|3 cooling (if there was no selectivity change, the correlation would be zero). We conclude that PIT multiunits lost selectivity across images as a function of RF location.

### Cooling did not reveal shape-specific deficits

We used regression analyses to find image features especially affected by cooling. These features could be anything encoded by PIT, including luminance, contrast^8^, orientation content, curvature content^9^ and categorical membership (e.g. “faces,” “body parts,” “tristars”, **Fig. 5**). In addition to these features, we used principal component analysis on our images to extract 87 different quantitative descriptors for each of our 293 images (see **Supplementary Fig. 5**). We also used experimental predictors, such as the mean population spike rate per image (before cooling), and the mean decoding accuracy evoked per image (before cooling). We used all 87 features in a linear regression analysis that could explain the change in decoding accuracy per image. In both monkeys, the only consistent predictor of V4-or V2|3-cooling accuracy loss was the magnitude of classification accuracy before deactivation: the larger the classification accuracy for each image before deactivation, the larger the subsequent reduction in accuracy. During V4 cooling, the percentage of variation explained by each model was 45-55% (monkeys R and G: R^2^ = 0.45 and 0.55, *P* = 4×10^−6^ and 1.74×10^−7^, F(100,246): 2.0, F(100,192): 2.4); during V2 cooling, the R^2^ values were 0.48 and 0.56 (*P* = 1×10^−7^ and 3×10^−8^, F(100,246): 2.3, F(100,192): 2.5). The linear model could not account for differences in decoding accuracy between V4 and V2|3 cooling (R^2^ = 29%, *P* = 0.84, 0.91, F(100,246): 0.8, F(100,192): 0.8). In summary, no shape-or category feature showed a statistical relationship with decoding accuracy reductions during cooling, suggesting that a yet-undiscovered image property is differentially represented among the pathways, or that image feature encoding does not differ between them. This oriented us to the possibility that the primary difference between these pathways is not between the bypass pathways, but between the long pathway and the bypass pathways themselves.

**Figure 5.**
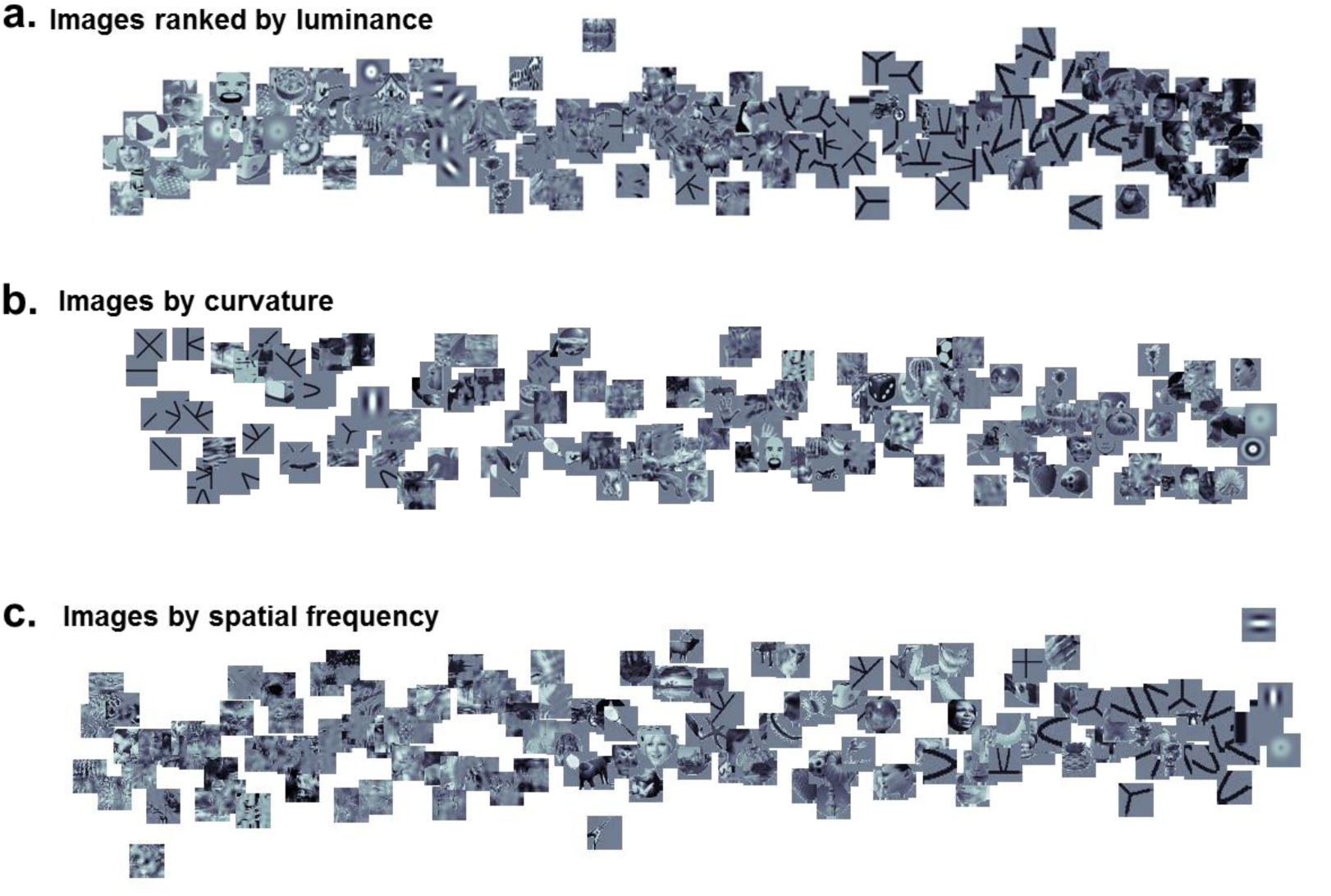
Examples of three image features used to predict changes in accuracy. **A.** Images sorted by their luminance value. **B.** Images sorted by their curvature content. **C.** Images sorted by the first Fourier-transform principal component (high vs. low spatial frequency content).

### Cooling parallels in the Standard Model of Visual Recognition

The previous results oriented us to compare PIT units that received inputs from the long (V1→V2→V4→PIT) pathway versus PIT units that received inputs from the short (V1→V2→PIT or V1→V4→PIT) pathways. This was outside our experimental reach, because both of our cooling interventions affected cells that depended on continuous information flow through the long pathway. Thus we proceeded by exploring simulated versions of these two PIT types, using the Standard Model of Visual Recognition, specifically the HMAX version of Serre, Oliva and Poggio (2007). This is a hierarchical, feedforward-only model inspired by the visual system^10,11^, comprising comprises multiple layers *(areas)*, each with many filters *(receptive fields)* of different sizes. The mean filter size per layer increases along the hierarchy, with small V1-like RF sizes in the first layer, and large AIT-like RF sizes in the last layer. Each layer performs two serial operations: a convolutional tuning operation followed by a pooling (invariance) operation. Inspired by the simple/complex cells in V1, the convolutional operation provides a measure of similarity between the input pattern and the “synaptic” weights of its filter. The outputs of different convolutional steps are then pooled using a maximum operation; this is a non-linear step that reduces multiple inputs into a single output, like a complex cell responding with its most active simple cell input. This pooling step results in fewer responses feeding into the next layer and more invariance to scale/position changes. At the highest layers of the model, there emerges a sparse population of units, whose activations encode an abstract representation of the original image. This response vector can be used in a final classification step to measure the accuracy of representation of the original image.

Our version of this model had three alternative pathways that could provide input to a given simulated PIT unit: the long pathway had four layers (V1→V2→V4→PIT); the second pathway skipped the second layer (V1→V4→PIT), and the third pathway skipped the third layer (V1→V2|3→PIT, **Fig. 6a**). Filter widths doubled at each layer, but for the bypass pathways, RF size quadrupled at the bypass stage. This insured that all filters in a given area had the same range of sizes, irrespective of their inputs. The key difference in filter shapes between the long-and short-pathway inputs was the level of detail present in the filters: for example, V4 units shaped from V2 inputs were more abstract than V4 units shaped from V1 inputs, because the latter set of filters sampled visual activity which had undergone one fewer round of max pooling. At the end of the network, we decoded responses using SVMs in an all vs. all approach, with leave-one-out cross-validation and shuffled-label control.

**Figure 6.**
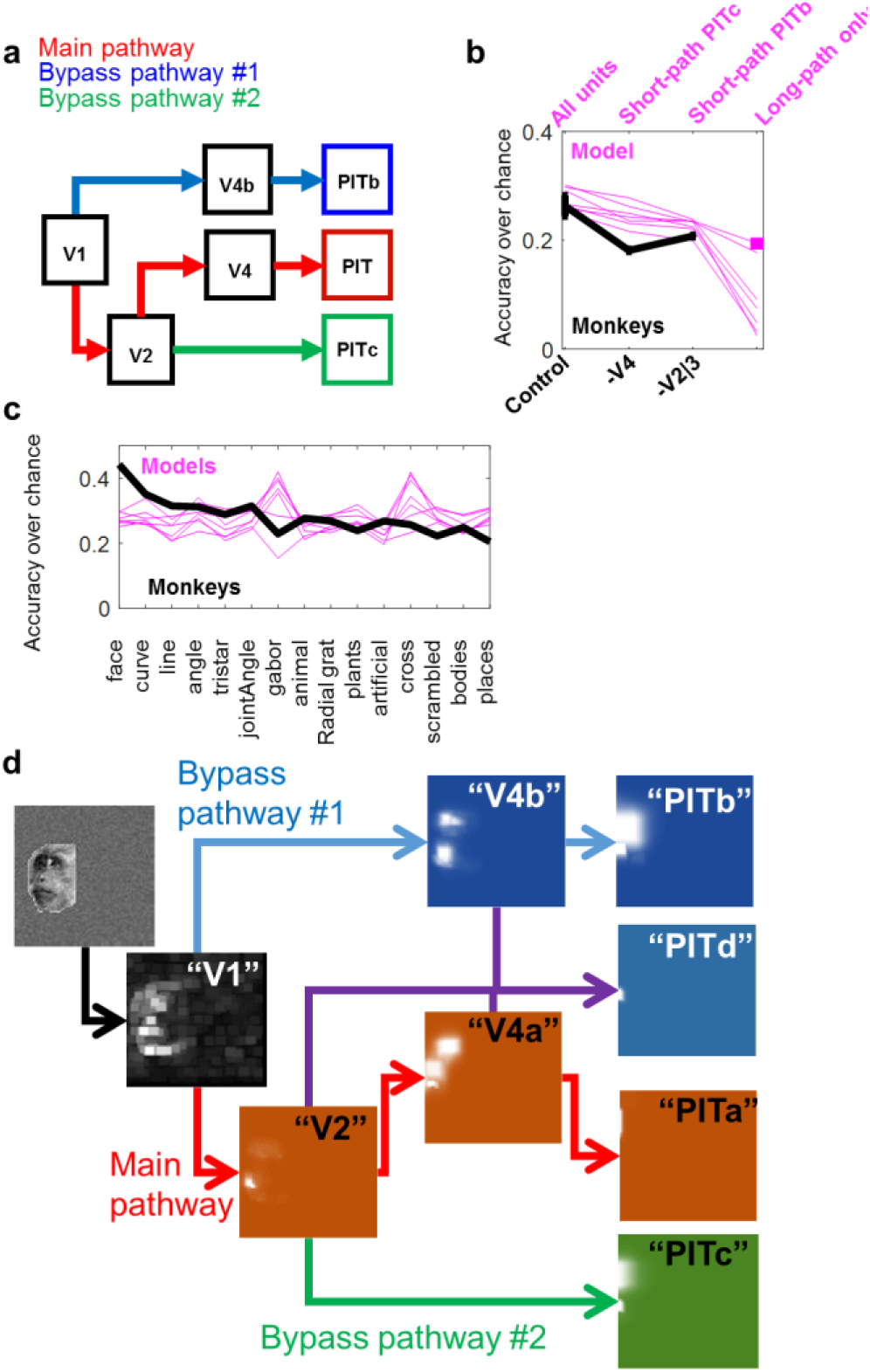
a. Modified standard model of visual recognition (HMAX architecture) **b.** Mean classification accuracy over chance (± SEM), for both animals, during the warm, V4 and V2|3 deactivation (black) and for all simulated PIT populations in seven models (pink lines). The label “All units” refers to mixed short-and long-pathway PIT cells. The square under the “Long-path only” label shows the accuracy achieved when units were pooled from seven networks, in order to match cell count. **c.** Accuracy per category for all seven models (pink). The black line shows the mean accuracy per category from the monkey data. **d.** Example of activation of units at different layers in the hierarchy.

There were 58 units total at the final layer: 4 long-pathway units, 25 V1→V4→PIT units, 25 V2→PIT units, and 4 units that received mixed inputs from long-and short-pathways. SVMs performed best at classifying each image when using a mix of long-and short-pathway units (N = 58), scoring 28±1% after baseline subtraction, mean of seven models ± SEM). For comparison, SVMs trained on the control monkey data showed an average of 26±2% after baseline subtraction. When SVMs were trained using data from only one population of simulated PIT cells, they performed worse than when using the mixed population (PITc: 24±1%, PITb: 24±1%, PIT: 9±3%; **Fig. 6b**). Within each model, the number of simulated PIT units at each short-pathway was the same (N = 25 units per final bypass layer), but the number of long-pathway units was lower because of the additional pooling stage (N = 4); this partially explains why SVMs performed so low when using long-pathway values only. To correct for this, we also trained SVMs using all the long-pathway units from seven different networks: pooled together, these 28 units still performed worse (19±1% above baseline) than SVMs trained on a mixed population. These reductions in simulated PIT cell diversity were similar to the accuracy from cooling data (mean −V4 deactivation performance was 16% and −V2|3 deactivation performance 19%, averaging both monkeys).

Unlike the monkey PIT units, which led to the highest classification accuracy for faces, the entire population of simulated PIT model units showed highest accuracy values for line shapes, bodies and artificial objects (24-34% over chance for all categories, **Fig 6c**). Visual examination of the activation patterns within each layer highlighted interesting differences (**Fig. 6d**). Units in the early layers (V2, V4) of the long pathway lit up local features, like eyes, but in subsequent layers (PIT) of the long pathway, these features were pooled into more complex feature selectivities. These long-pathway units looked like they would be good at discriminating complex images, but at the expense of representing more primitive geometric features. In contrast, PIT units receiving inputs from the bypass pathways seemed to preserve the explicit representation of the more primitive sub features. We tested this observation as follows. We compared the image selectivity to local and global features of two classes of simulated PIT cells - those at the terminus of the long pathway or the short V1→V4→PIT pathway. We used images of 10 faces and 10 quadrupeds from our 293-image set. We used SVMs to interrogate the long-and short-pathway units on four tasks, where success in each task depended on discriminating local vs. global features. The tasks were to classify 1) faces vs. quadrupeds, 2) heads vs. faceless heads, 3) faces vs eyeless faces, and 4) quadrupeds vs. legless quadrupeds (**Fig. 7**). The first two comparisons involved many local features (a global comparison), the second two comparisons involved a local feature. The pathways performed differently: for the global task, the long-pathway units showed better performance than the short-pathway units (accuracy classifying faces vs. heads, long-pathway: 0.58±0.05, short-pathway: 0.54±0.06; faces vs quadrupeds: long-pathway: 0.58±0.05, short-pathway: 0.57±0.07). For the local-feature tasks, the short-pathway units allowed better performance than the long-pathway units (accuracy detecting missing eyes, long pathway: 0.58±0.05, short pathway: 0.64±0.04; accuracy detecting missing legs: long pathway: 0.53±0.06, short pathway: 0.66±0.06). This suggested that in the macaque brain, short pathways could be helpful for fine, local-feature discriminations, while the long pathways could implement Gestalt-like, global discriminations.

**Figure 7.**
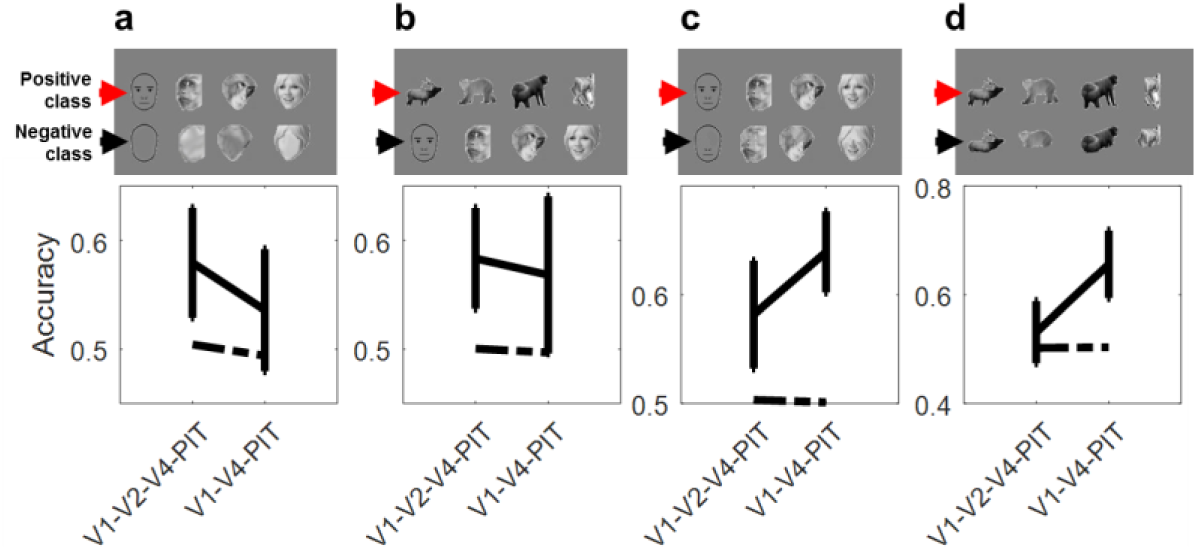
Sensitivity to image parts by long-and short-pathway units. Classification accuracy differences among different pathways. Solid lines show mean SVM accuracy values, error bars show standard error of the mean. Broken lines show accuracy after label shuffling. **a.** Heads vs. faceless heads. **b.** Faces vs. quadrupeds. **c.** Faces vs. eyeless faces. **d.** Quadrupeds vs. legless quadrupeds.

### Cooling affected fine discrimination less than coarse discrimination

The cooling interventions always disrupted the long pathway but preserved at least one short pathway. If the short pathways are more important for local (fine) discriminations and the long pathway more important for global discriminations, then during cooling, SVMs should perform better with local discriminations because of these remaining short pathways. We wanted to test this hypothesis, so first we had to define “local” vs. “global” discriminations in the context of our data.

The decoding accuracy analysis (**Fig. 2**) showed that monkey PIT populations discriminated some images more accurately than others. The image pairs that were hard to discriminate must have had similar locations in the neural activity space: the closer the images, the more likely that individual trials were on the wrong side of the dividing linear hyperplane. We ranked the decoding accuracy for each image as a function of the Euclidean distance to its neighbors (in the original 100+ dimensional activity space). This plot showed that decoding accuracy for two images increased with their distance in activity space. It also showed that very close neighbors were systematically misclassified (**Fig. 8a**, bottom). Thus we can interpret distance as a continuous metric for local-feature vs. global-feature discriminations: close image neighbors activate similar feature detectors in PIT relative to distant image pairs.

**Figure 8.**
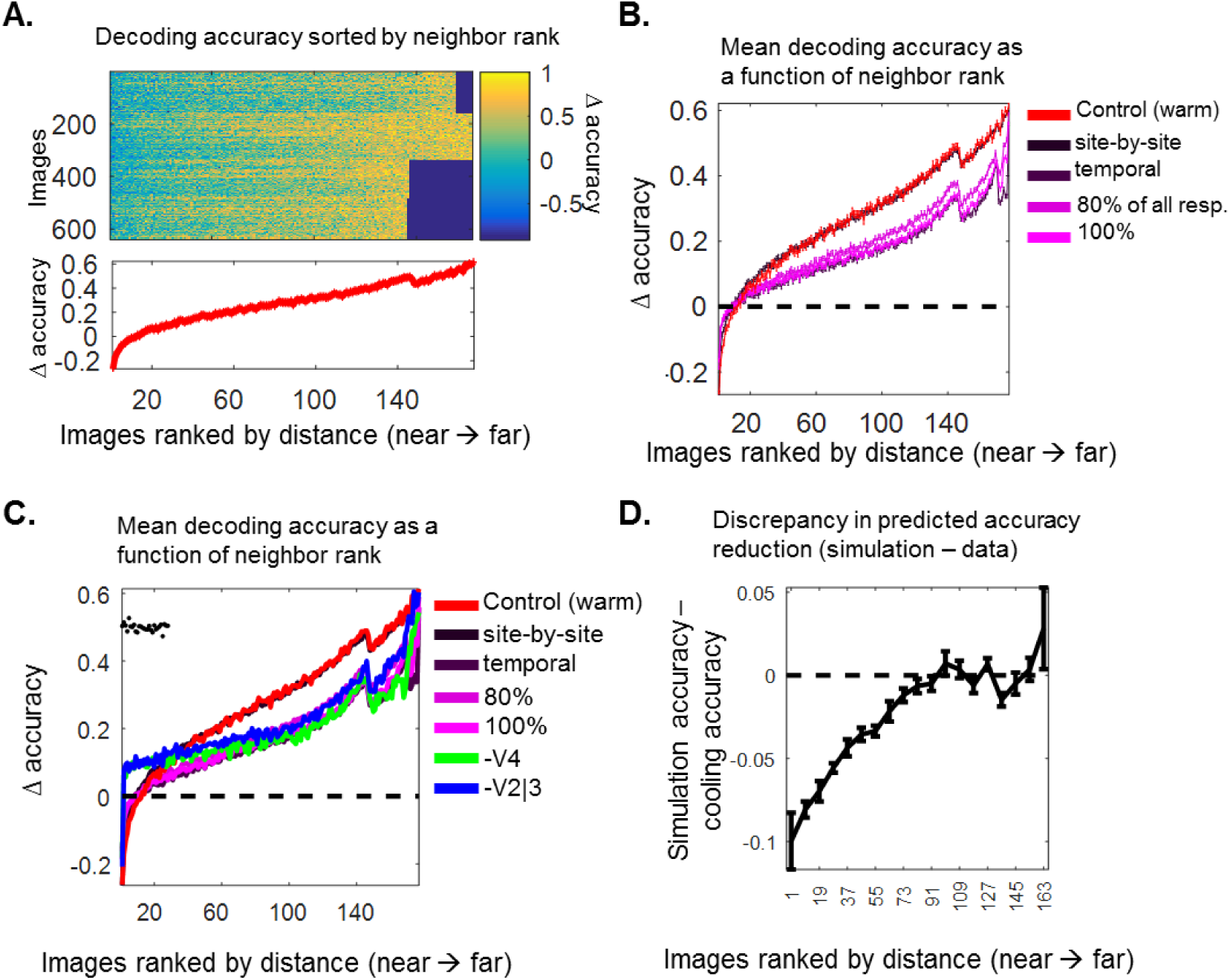
Differences in decoding accuracy for near-vs. far neighbors. **a.** Linear decoding accuracy for every image (rows) when compared to its neighbors, as sorted by rank distance (columns). Top figure shows the decoding accuracy for all images, across days and monkeys. Bottom figure shows the mean decoding accuracy for each rank position (±SEM). Colors show the decoding accuracy minus the shuffled-label accuracy. **b.** Mean decoding accuracy (minus shuffled baseline, ± SEM) measured before cooling (red). The additional curves, colored in magenta, show the values for the different cooling simulations (see Figure 3 for a description of each mechanism). **c.** Same as (c), but with data from V4 cooling (green) and V2|3 cooling (blue). Black dots show which image ranks show statistically differences between the cooling data and trial-by-trial model. **d.** Discrepancy in predicted accuracy reduction between the simulation (trial-by-trial) and V4 cooling data. Each point shows the average for 10 neighbors (±SEM), from nearest to furthest.

We then asked whether the cooling reduction in accuracy varied with neighbor distance, and if so, whether that change could be predicted by a random cooling mechanism like those of **Figure 3**. These random cooling mechanisms were blind to the identity of each image and to their distances in activity space. This created two hypotheses: the first hypothesis was that the change in decoding accuracy as a function of distance will be either a simple multiplicative or subtractive change, captured by random cooling mechanisms. The second hypothesis is that the change in decoding accuracy as a function of distance will not be explained by a simple gain change, but in fact will be better preserved within short distances, as predicted by the convolutional model of visual recognition.

We plotted the decoding accuracy as a function of distance for the random cooling mechanisms and found that they showed a series of multiplicative changes to the control (warm) accuracy-distance plot (**Fig. 8b**); these curves showed small changes in accuracy for near-neighbors, and larger accuracy changes for far neighbors. In contrast, when we plotted the cooling data’s decoding accuracy as a function of distance, we found that these curves differed from the simulated-cooling curves within the range of short distances (**Fig. 8c-d**). The random cooling mechanism curves predicted small decrements in misclassification, but the cooling data showed a considerable improvement in classification. The cooling data and the simulated cooling data curves were otherwise well-matched at the mid-to far range of neighbor distance. To compute the statistical reliability of this observation, we derived the probability that the cooling median accuracy at each distance could be emitted by the simulation (we ran a Wilcoxon rank-sum test at the *i*th neighbor position, asking if the median accuracy measured in the cooling distribution was higher than the median accuracy derived from the simulation). We found that the nearest 28 neighbors were reliably better classified than predicted by the simulation (*P* values ranged from 4×10^−35^ to 0.03 within the nearest 28 positions, one-tailed paired Wilcoxon signed rank test, N = 640 images, Z-values = 1.9-12.3). This resilience in decoding accuracy for near neighbors is consistent with the interpretation that short pathways are most helpful with fine discrimination.

## DISCUSSION

We investigated how posterior inferotemporal cortex cells combine information from areas V2, V3 and V4. We implanted microelectrode arrays in PIT while cooling areas V2/V3 or area V4. PIT multiunits showed a reduction in firing rate that was similar across both types of interventions. Support vector machines showed that classification accuracy was reduced more during V4 deactivation compared to V2|3 deactivation. Changes in classification accuracy could not be predicted by any class of visual features, such as contrast or spatial frequency content. This finding is consistent with previous iterations of the Standard Model of Visual Recognition, which implemented a hierarchical convolutional network where different bypass projections contain the same types of visual features. This previous model also modeled bypass pathways and predicted a role for fine discrimination, and here we explore that idea further^2^. We confirmed that these pathways provide an advantage to simulated PIT units in fine discriminations. According to the model, the key contrast was between long-pathway PIT cells (V1→V2→V4→PIT) vs. short-pathway PIT cells (V1→V4→PIT), and these populations could not be isolated using our cooling technique. However, we found that images that are represented similarly in PIT activity space were better discriminated during cooling; this advantage was not predicted by non-specific cooling simulations.

Our principal conclusion is that input pathways of different lengths create a diversity of functional preferences in a given cortical region. We argue that multiple pathways allow some units to respond to complex features, and some units to simpler features. This raises several questions. First, in the context of a fine discrimination task, why would units with complex image preferences be less helpful than units with simpler image preferences? Cells with complex image preferences can be less responsive to variations of their preferred image: Kobatake and Tanaka have shown that IT cells have a minimum number of “critical features,” a minimal combination of image parts required to elicit a response^12^. If any of those features are missing, the cell may not respond at all. Thus in the context of detecting the presence of simple parts within a complex image, there may be an advantage of having cells tuned for the complex image along with cells tuned for its simple parts. This type of simpler preference is more likely to arise earlier in the visual hierarchy and become preserved by short bypass pathways. Did we miss any classes of 2-D, luminance shape features that might be differentially represented between the V2|3 and V4 paths? While possible, there are no strong theoretical candidates for such features. Hegde and Van Essen (2007) compared the relative shape selectivities in neurons from V1, V2 and V4, and showed that these cells responded similarly to the same set of simplex and complex images, offering little qualitative diversity^13^.

The V1−V2|3−V4 input pathway is the most important source of excitatory activity to PIT, and one might expect that its disruption would extinguish nearly *all* activity within the PIT scotoma. However, PIT units within the scotoma continued to respond, albeit weakly. This was because we did not cool all of V2, V3 or V4 - the scotomas covered only a few degrees of central vision, and near-foveal representations receive a disproportionate amount of cortical real estate in the brain. Indeed, cortical areas deep in the visual hierarchy receive too many inputs and may be practically impossible to silence without bilateral V1 resection, silencing of lateral connections and feedback: as an extreme example, anterior IT cells show no statistical change in overall firing rate after surgical resection of areas V4 and PIT^14^. Ultimately, we will get more answers to the problem of multiple pathways in macaque by recording from single units with known anatomical input profiles, a goal that may be met with the use of chemo-or optogenetics.

## METHODS

All procedures were approved by the Harvard Medical School Institutional Animal Care and Use Committee, following the Guide for the Care and Use of Laboratory Animals (8th edition, National Academies Press). This paper conforms to the ARRIVE Guidelines checklist.

### Behavior

Two adult male macaques (10 −17 kg) from the New England Primate Center were trained to perform a fixation task. The task required them to stare at a 0.5°-wide red or black square in the middle of the screen, keeping their gaze within ±1.3° from the fixation spot. We used an ISCAN eye monitoring system to keep track of eye movements (www.iscanin.com). The trial timeline was as follows: at the start of each trial, the fixation target appeared and the animal had up to eight seconds to direct its gaze to the fixation target. Once fixation was acquired, a small reward could be delivered to encourage the animal. Within a random period between 17- 117 ms after fixation onset, an image appeared perifoveally for 200 ms, then disappeared for 200 ms until a new image appeared. This on-off cycle could be repeated with 3-5 different images per trial. If the animal held fixation until the end of the final on-off cycle, a reward was dispensed. The reward size increased by 25% of the initial reward size every 100 trials.

### Animal care

The animals were housed in a vivarium with a 12-h light/dark cycle, under social pairing. They worked during the daytime. The monkeys had no previous major surgical history.

### Visual stimuli

We used MonkeyLogic to control experimental workflow (http://www.brown.edu/Research/monkeylogic/). We used 293 high-resolution images, including photographs and simple shapes. Photographs came from Google Images, and our choice of pictures were guided by categories used in previous IT studies^15,16^ including animals, artificial gadgets, body parts, faces, places, plants/fruits(20-21 examples per category; faces and body parts were evenly distributed between monkey and human). Most of these images were used to create scrambled counterparts via the Portilla and Simoncelli visual texture model^17^, which transforms white noise into textures that share pairwise joint statistical constraints as the original intact images. These textures convincingly replicate small shape primitives present in the original images and scatters them throughout the image (118 textures). We also used 54 simpler line shapes such as lines, curves, tristars, radial and linear gabors, and simple combinations of lines and curves (joint angles). These line shapes were generated using the Cogent Matlab toolbox (developed by John Romaya at the LON at the Wellcome Department of Imaging Neuroscience, http://www.vislab.ucl.ac.uk/cogent_graphics.php). Images were 1.4° in width (for monkey G) or 2.0° (for monkey R) at their longest axis. The images were not normalized for luminance, contrast or other visual properties.

### Implanted devices

The cryoloops were manufactured in the laboratory of Stephen Lomber and are described in Lomber, Payne and Horel (1999)^18^. Cryoloops were composed of 23-gauge hypodermic stainless steel tubing, shaped to fit the individual curvature of each animal’s occipitotemporal shapes as determined by structural magnetic resonance images. The cryoloops were 3.5 mm wide and between 4-11 mm long. A microthermocouple sensor was attached to the stem of the cryoloop to monitor its temperature. The bodies of the cryoloops were wrapped in Teflon tubing except at the loop. The loops contained protected inlet/outlet ports that permitted the daily connection of Teflon tubes carrying chilled methanol, as driven by FMI “Q” Pumps (Model QG150, fluidmetering.com). The methanol was contained within the tubing system and could not cause any chemical harm to the tissue. The custom floating microelectrode arrays were manufactured by MicroProbes for Life Sciences (Gaithersburg, MD); each had 32 platinum/iridium electrodes per ceramic base, electrode lengths of 4-16 mm, impedances between 0.7- 1.0 MO, all connected to a 36-channel Omnetics connector (allowing for two additional grounds and two reference electrodes).

### Surgical procedures

Both animals were implanted with custom-made titanium headposts before fixation training. After several weeks of post-surgical recovery and fixation training, the animals underwent a second surgery for the implantation of cryoloops and floating microelectrode arrays. Animals were anesthetized using ketamine/xylazine (I.M.) and isoflurane; buprenorphine/non-steroidal anti-inflammatories were used for pain control. In each animal, we performed a craniotomy centered at the lunate sulcus and extending antero-laterally. Monkey R received three cryoloops, two placed within the left lunate sulcus and one over the prelunate gyrus. The medial lunate sulcus loop was located 20 mm from the midline, traveled 7 mm deep into the sulcus and was 3 mm wide; the lateral lunate sulcus loop traveled 4.5 mm into the sulcus and was 3.5 mm wide); the prelunate gyrus loop was placed anteriorly to the lunate sulcus loops, was 11 mm long and 3 mm wide. Monkey G received two cryoloops, one over the prelunate gyrus and one within the lunate sulcus. The lunate sulcus loop was placed 2.1 cm from the midline, was 11 mm long with this axis running in the mediolateral axis within the lunate sulcus, 3 mm wide, and its most dorsal edge was 1.5 mm deep. The prelunate gyrus loop was also placed 2.1 cm from the midline, anteriorly to the lunate sulcus loop, ran 10.5 mm long and was 3 mm wide. We collected thermal images to map the spread of cooling from the tubing, and confirmed that it was limited to 1-3 mm radially, as first shown in previous publications^19^. Two to three floating microelectrode arrays were implanted within the same intraoperative session, after placement of the cryoloops. Their insertion sites were determined using three guidelines: they had to be anterior to the inferior occipital sulcus, many millimeters away from the prelunate gyrus cryoloop, and avoid large vasculature. All arrays were implanted caudal to the posterior middle temporal sulcus. We implanted two 32-channel arrays in monkey R, and one 32-channel plus two 16-channel arrays in monkey G.

### Experimental session workflow

All data reported was collected within a couple of months after implantation. Each day, the animal would be head-fixed and its implants connected to the experimental rig: first the cryoloops were connected to the chilled-methanol-bath tubing and temperature sensors, then the microelectrode arrays were attached to their headstages. The first step each day was to calibrate our measurements of the animal’s gaze using the built-in MonkeyLogic routine. We used the Plexon Multichannel Acquisition Processor (MAP) Data Acquisition System to collect electrophysiological information, including high-frequency (“spike”) events, local field potentials and other experimental variables, such as eye position, reward rate and photodiode outputs tracking monitor frame display timing. Each channel was auto-configured daily for the optimal gain and threshold; we collected all electrical events that crossed a threshold of 2.5 standard deviations from the mean peak height of the distribution of electrical signal amplitudes per channel. These signals included typical single-unit waveforms, multi-unit waveform bursts, and visually-active hash.

The animal began its fixation task while we collected responses from the arrays with the cryoloops at body temperature (36-37°C; this is what we call “control” or the “warm” condition). After ~20 minutes of data collection to permit ~4-6 repetitions of each image, we activated either the V2|3 or V4 cryoloops, bringing the temperature of the cryoloops to ~9°C, which lowered the temperature of the adjacent cortex to 16-18°C. We waited for another 5 repetitions of each image to pass, and then turned off the cryoloop pumps and collected 1-2 more repetitions under this first re-warming session. We then paused the fixation task for 10 minutes to allow the tissue temperature to return to normal and preserve the animal’s motivation for a second round of cooling. After 10 minutes, the temperature reported by the cryoloops was around 34°C, and we re-started our experiment. We repeated each image presentation 3-4 times and then activated the second set of cryoloop(s) (~8°C), waited for 5 repetitions and turned off the cryoloops. We then collected data until the animal was satiated. We balanced the order of the V2|3 vs. V4 cryoloop activations: if on the first day we activated the V4 cryoloop first and V2|3 cryoloops second, the next day we activated the V2|3 cryoloops first and V4 cryoloop second. There was an even number of days for each cooling order.

### Electrophysiology data preparation

The raw data files comprised event (“spike”) times per channel for the entire experimental session (the number of channels available per day were 64, but not all provided reliable signal-to-noise qualities). We divided each daily data set into thousands of raster plots defined by the onset of each image presentation and labeled each raster plot with its corresponding channel, image name and temperature condition. We defined three windows of analysis: the baseline period lasted from 0-50 ms after image onset, the early period from 51-150 ms after image onset, the late period from 151-250 ms after image onset, a full image presentation window was 51-400 ms after image onset. We found that multiunit responses could last almost 400 ms, although their peak responses always occurred within the early window. Here we report responses within the full window minus the activity within the baseline window (we call these evoked responses). For all multivariate analyses, we normalized the activity of each site by transforming its evoked responses to z-scores: all evoked responses emitted by a single site during an experimental daily session were averaged, this mean response was subtracted from all individual evoked rates, and each value was then divided by the standard deviation of all evoked responses.

Although our full dataset contained 293 images, we did not have enough time to present all images every day and still get the minimum of 15 presentations across the control and cooling conditions. Thus we presented over half of the total image set each day (10 images from each complex category, such as faces and places, along with half of the scrambled textures per day, with most of the simple line shapes, rounding to about ~170-177 unique images per day). The responses of a given channel were correlated across days, but were also statistically different by multivariate descriptors such as multi-dimensional scaling. Because of these differences, we did not combine channel information across days and instead created a multi-day pseudo-population, where sets of concurrently recorded channels (*N* = 50-64) from different days were treated as if recorded at the same time (see Ch. 19 of reference 20)^20^. Thus the final activity space is defined by firing rates collected across “site-days,” where some dimensions represent responses from the same channel to the same image collected on different days. Because the whole image set was presented on different days, we had two pseudopopulations per animal, each containing different site-day responses to each half of the image set. Each of our pseudopopulations had between 100-300 multi-units.

### Scotoma mapping experiments

The goal of these experiments was to identify the parts of the retinotopic field that were captured by our arrays, and the relative location of the response impairment caused by cooling. To achieve this goal, we had the animals fixate while we presented a single image (black-and-gray cartoon face, 2.0°-wide) within all positions in a radial grid (angular coverage of 0-315°, 45° steps; radial coverage of 0-8° from the center of the screen, in 0.5° steps). Three to five positions were randomly chosen per trial. After data collection, we defined evoked responses per position as follows: first we quantified the firing rate per site during the early window of activity (51-151 ms after stimulus onset) and then subtracted the firing rate per site during the baseline window of activity (0-50 ms after stimulus onset). We averaged these evoked responses per position within each site and used the griddata.m Matlab function to interpolate the scattered data into a continuous map. This map was smoothed using a 1°-diameter disk filter. This map represented the aggregate receptive field of each multiunit site in our arrays. To identify the overall scotoma, we averaged the response fields of all sites during the control condition and subtracted the average response fields of all sites during V2|3 or V4 deactivation. We measured the size of each scotoma by hand, using the calcArea.m function (http://www.mathworks.com/matlabcentral).

### Firing rate and latency

The goal of these analyses was to measure changes in the overall firing rate (excitatory drive) of PIT multiunits during input deactivation. These changes included the amplitude of peristimulus rate histograms (PSTHs) and the latency of response. To quantify the changes in evoked response magnitude, we computed the evoked responses per site as described in *Ephys Data Preparation* and averaged these responses across all channels within each temperature condition. We did the same operation using z-scores. We calculated the probability that the median responses emitted during each temperature condition (control, V4 and V2|3 cooling) were sampled from the same distribution using a Kruskal-Wallis one-way analysis of variance. To determine if there was a statistical difference between the V4 and V2|3 cooling condition responses, we used the Wilcoxon signed rank test for zero median. For the latency analyses, we obtained the mean PSTH in response to each image, per site and temperature, and then stacked all image-specific PSTHs in a matrix measuring N_images_ × 400 (ms after stimulus onset). We identified the time when each PSTH exceeded two standard deviations over baseline and called this *response latency*, with the only acceptance criteria that a plausible response latency would only occur between 30-200 ms after image onset. We also computed the earliest time point when all PSTHs demonstrated the greatest variance in amplitude, as an indicator of the *tuning* latency.

### How we identified channels with reliable visually driven activity

Many electrodes in the arrays reported electrical activity that was not visually driven, possibly because the electrodes were on the pial surface. We repeated this analysis only using channels that showed a statistical difference in mean activity between the baseline and evoked time periods. Using a cross-validation approach, we used 5% of all trials to perform a Wilcoxon signed rank test for the median rate difference during each interval. This told us which channels showed a statistical difference in rate during visual stimulation. We then used the remaining 95% of trials to compute the firing rates during baseline and evoked windows for the selected channels. Monkey R’s arrays showed 38 out of 64 visually responsive sites; monkey G, 30 out of 64 (*P* < 0.05, two-tailed Wilcoxon signed ranked test for zero median).

### Encoding accuracy analyses

We trained support vector machines with a linear kernel using the Matlab function fitcsvm.m. We used an all-vs.-all approach, with SVMs trained to discriminate between pairs of images, using leave-one-out cross-validation. There were 4-5 response vectors per class within each comparison (the data used for classification were Z-score vectors; see *Spike Data Preparation*). To estimate the chance accuracy for each paired comparison, we concurrently trained SVMs using the same set of data vectors but with shuffled labels. The number of vectors for each two-class comparison was small, and thus we found that chance accuracy values could vary between 0-1 across all comparisons; the median shuffled-label misclassification rates were 0.60-0.63 for monkeys R and G. We subtracted the chance, shuffled-label accuracy classification rate from the correct-label accuracy classification rates to account for this bias. As an insight to explain this deviation from the expected chance accuracy of 0.5, we trained SVMs to distinguish between stimulus categories (listed in the *Visual Stimuli* section). Each category pair comparison involved 10-20 times as many response vectors as the individual image-vs-image SVM analyses, and the dataset was otherwise identical. Here we found a more reassuring shuffled-label statistical baseline of 0.50 in both animals. Both the category and image-per-image SVM accuracy analyses led to the same conclusions presented in the *Results* section.

### Projection analysis

The goal of this analysis was to reconcile the findings that cooling V2|3 and V4 led to equivalent reductions in PIT population firing rates, but different reductions in classification accuracy. We measured the cooling trajectory traveled by each image during V2|3 or V4 cooling and to project it onto the direction of a minimum response vector, where this direction represented a non-specific reduction in firing rate across all sites. We conducted the analysis as follows; first, for every pseudo-population, we defined a minimum response vector **v**_**min**_ = {min arg(**x_j_**)} = {min arg(*x*_*1*_), min arg(*x*_*2*_), … , min arg(*x*_*n*_)}, where ***x***_**j**_ is a variable representing all the mean responses from the *j*th array site and *n* is the number of sites, thus the vector length is *n*. Next, we computed three mean vectors per image: **v_warm_** = mean response vector for the given image during the control condition, also of length *n;* **v_-v4_** = mean response vector for the given image during V4 cooling and **v_-v2_** = mean response vector for the given image during V2|3 cooling. We computed the cooling trajectory vector of each image as **v_warm-v4_** = **v_warm_** – **v_-v4_** and **v_warm-v2_**= **v_warm_** – **v_-v2_**. Finally, each individual trajectory was projected onto the vector **v_warm_ - v_min_**; the projected (parallel) component was subtracted from the cooling trajectory vector to compute the perpendicular component.

### Cooling simulations using the control data

We simulated the effects of cooling on the control decoding accuracy, by applying fractional reductions to the control spike rates. The approach was to first identify the fractional reductions in firing rate for all channels during cooling in a given pseudo-population and then to use this fractional distribution to simulate cooling changes with data from a different pseudo-population. Fractional reductions were defined as *ƒ*_*i*_ = *r*_*i*_, _*cooling*_ / *r*_*i*_, _*control*_, where *r*_*i*_ is the mean firing rate for a given channel *i* in a pseudo-population *i* = 1-300. We sampled values from each fractional distribution with replacement and multiplied warm firing rates in three different ways. In all cases, we can envision the control data set as a matrix of dimensions *r × c*, where rows are multiunit sites (channels) and columns are individual image presentations. In the first mechanism, *site-by-site cooling*, we sampled *r* fractional changes and multiplied each sample times all the responses in one channel. In the second mechanism, *temporal cooling*, we sampled *c* fractional changes and multiplied each fractional change times all elements in the column. Finally, in the third mechanism, we randomly selected a given percentage of responses in the matrix, and multiplied them by an equal number of sampled fractional values (the *mixed site-by-site* and *temporal cooling*). We used each transformed control matrix to train and test support vector machines as described above.

### Selectivity analyses (F-statistics)

In this analysis, the F-statistic was used as a measure of selectivity for each multiunit. The F-statistic is a ratio of mean squares, specifically the mean square error estimate for the variance of responses among images, divided by the mean square error estimate for the variance within each image. We computed each F-statistic in a channel by channel basis using the responses to all images within each temperature condition. For each channel, one F-statistic was computed using the warm data (*F*_*control*_), another using the V4 cooling data (*F*_−*V*4_) and another using the V2|3 cooling data (*F*_−*V*2|3_). We plotted each *F*_*control*_ against its paired *F*_−*V*4_ and *F*_−*V*2|3_ values. To determine if the slope in each given scatterplot was different from unity, we used a bootstrap approach, where we computed 1,000 different slopes by sampling each channel with replacement (note that we kept each F-ratio trio together; we did not mix warm and cooling F-statistics from different channels). We then asked if the slope distribution from this bootstrap included 1.

We used randomization to measure any differences between the mean slopes computed during the V4 and V2|3 cooling conditions (that is, whether there was a difference between the mean *F*_−*V*4_ / *F*_*control*_ slope vs. the mean *F*_−*V*2|3_ / *F*_*control*_ slope). The null hypothesis is that the mean V4 and V2|3 slopes came from the same distribution. Therefore, we had to create this null distribution. In each of 1,000 passes, we randomly mixed the labels between the V4 and V2|3 F-statistics for each channel and computed a *F*_*cooling*_ / *F*_*control*_ slope. We did that twice per pass, and then subtracted the two slopes. After 1,000 passes, we had 1,000 slope differences that we then compared to the experimental slope difference. We found that these null difference distributions were defined by 5^th^ and 95^th^ percentile values of −0.07 to 0.07 (monkey R) and −0.05 to 0.05 (monkey G). The observed differences in mean cooling slopes were −0.18 and −0.06 (monkeys R and G). The probability that the experimental differences in V4 and V2|3 slopes came from such mixed distributions were 0.001 and 0.02.

### How we defined response field overlap with the scotoma, for the *F*-statistic analysis

For each channel, its RF overlap was defined as the average number of spikes emitted in response to stimuli presented in the stimulus/scotoma region, divided by the total number of spikes emitted in the central 8×8°. The mean RF overlap value was 0.12±0.01 and 0.15±0.01 (monkeys R, G).

### Linear regression model

The goal of this analysis was to determine whether the change in classification accuracy during V4 cooling or during V2|3 cooling could be predicted using different image features. The regression matrix had dimensions of 293 × 87 (images × visual features). The features were luminance (defined as the mean pixel value transformed by the monitor’s gamma function), contrast (variance of the pixel values transformed by the monitor’s gamma function), horizontal vs. vertical power (obtained via a wavelet decomposition analysis using the Matlab function wavedec2.m), curvature (defined by the variance of each image’s discrete Fourier transform spectral power around all orientations), 50-pixel-based principal components as defined by the pca.m function), 30 spatial frequency principal components (pca.m applied to the discrete Fourier transformed images), categorical membership (defined *a priori* as angles, animal, artificial, bodies, cross, curve, face, gabors, radial gabors, joint angles, line, places, plants, scrambled, tristar), the mean population control firing rate per image and control classification accuracy per image. Values within each feature group were z-scored before fitting. The dependent variables were either 1) accuracy loss during V4 cooling (control accuracy per image minus −V4 accuracy), 2) accuracy loss during V2|3 cooling (control accuracy per image minus −V2|3 accuracy) or 3) the difference in accuracy loss during V4 minus V2|3 cooling ([control accuracy per image minus −V4 accuracy] - [control accuracy per image minus −V2|3 accuracy]). The probability that the linear model differed from the constant model was obtained two ways: first, we used the t-statistic provided by the “fitglm.m” function; second, we used a randomization test where the dependent variable was fit with a regression table made up of random numbers, sampled from a flat distribution. The table had the same dimensions as the true data matrix table. The R^2^ values of a thousand randomization tests were compared to the R^2^ from the regular regression table. To identify the most interesting predictors, we looked at all 87 regression weights and their t-statistics. There was one clear outlier: the control classification accuracy (t-statistics 8.6-8.7). We also fitted a regression analysis that penalized the number of regression weights (the Lasso). To insure that this was not simple regression to the mean, we also divided our control trials such that the control classification tuning curve used for regression was not the same as the control classification tuning curve used to calculate the cooling difference in classification accuracy (the estimated correlation between these cross-validated data sets were 0.33-0.45 for monkeys R and G, *P* < 10^−8^, two-tailed Student’s T-test).

### Standard model of visual recognition

Our computational model was based on an implementation by Serre, Oliva and Poggio (2007)^5^, available at http://cbcl.mit.edu/software-datasets/standardmodel/index.html. This model belongs to the family of hierarchical feedforward models (HMAX) by Riesenhuber and Poggio (1999)^21^ and developed over subsequent publications^2,22,23^. The model represents the visual object recognition system as a series of convolutional and pooling operations, which transform an image from pixels into neuronal responses. These responses can be used in a statistical classifier to decode their abstracted representational content.

The architecture of our network was three to four layers deep and contained three pathways: one main pathway and two bypass pathways. The main pathway had four layers: layer 1 (representing V1), layer 2 (V2|3), layer 3 (V4), and layer 4 (PIT units receiving inputs from the main pathway). The second pathway had three layers: layer V1, layer V4b (representing units in V4 receiving direct input from V1) and layer PITb (PIT units receiving input solely from the V1→V4 inputs). The third pathway also had three layers: layer V1, layer V2|3 and layer PITc (units in PIT receiving input directly from V2|3). These three types of “PIT” neurons showed different kinds of activation patterns, which we could decode using support vector machines.

#### Layers

Each layer represented a stereotypical set of operations: a convolution/tuning operation and a pair of max operations. The tuning operation is equivalent to a simple cell, which convolves the input with a filter bank via the tuning function 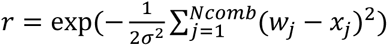, where *σ* = sharpness parameter, N_comb_ is the number of filters to combine, w = filter weight and x = input image or activity. This simple cell operation describes the Euclidean distance between the RF shape and the incoming input. Different simple cells are characterized by different shapes and sizes of their RF patches. There are more than one receptive field sizes at each layer, and each filter-size convolution is performed in parallel. Several outputs of this tuning operation are then combined in a complex-cell-like operation. *Complex cells* perform a pooling operation: they receive inputs from *N*_*s*_ simple cells with different RF sizes and compute the maximum response emitted by the set. Thus the output of a complex cell layer is sparser than the output of a simple cell layer, because maximum values are repeated across limited areas of response space. These complex layer responses are finally subsampled, imitating the decreasing number of cells that can cover visual space as one moves down the visual pathway.

Building the model required two major implementation stages: first we had to create receptive field (RF) patches for each layer and second, use these RF patches to compute responses to our experimental images. As in the 2007 publication, the patches were imprinted using experience-dependent activity.

#### Filters

Each layer contained a set of up to 200 unique filters: the V1 layer filters were Gabors at four orientations and eight sizes (3-10 pixels wide, or 0.1-0.4° wide given our monitor distance). Subsequent layer filters were imprinted using random samples of activity from the preceding layer. To train these filters, we randomly selected images from the Caltech-256 database and passed them through the V1 layer: the resulting activity patterns were sampled randomly to imprint filter shapes for the V2 layer^22^. Another set of images was passed through the V1→V2 layers, and that set of activity patterns was used to imprint filter shapes for the V4 layer. We did this for every layer in the long and short pathways. After repeating this process hundreds of times, this resulted in a model with different filter shapes at each layer. Filter sizes doubled at every hierarchical step, with the exception of the bypass pathways, where filter sizes quadrupled in width at the one skip level (e.g. V4 filters in the V1→V4 pathway were four times the size of V1 filters; PIT filters in the V1→V2|3 pathway were four times the size of V2|3 filters). To make sure that the RF shapes would match the statistics of natural images presented close to and far from the fovea, we also imprinted using differently sized variants of the same images (1°-, 2°- and 4°-wide versions of the same image).

#### Experimental image testing

We began with 293 different images for our experimental dataset. In the electrophysiology experiments, we presented each image multiple times and obtained a distribution of correlated but non-identical response vectors for each individual image. To mimic this trial-by-trial variability in the model, we created six variations of all 293 images, adding pixel-by-pixel noise and random changes in position, simulating differences in fixational eye movements. We then transformed our 293×6 experimental images into simulated PIT responses using the fully assembled network, and used SVMs to measure the classification accuracy from the model PIT units, SVMs were used in an all-vs.-all approach, with leave-one-out cross-validation and shuffled-label randomization control. The key theoretical contrasts were the relative performances between PIT units receiving inputs from different pathways: the V1→V2|3→V4→PIT pathway, the V1→V4→PIT pathway and the V1→V2|3→PIT pathway. We considered the long-pathway PIT units to represent our control temperature population, the latter the deactivated state units.

#### Local vs. global feature analysis

The purpose of this analysis was to determine whether closely represented images suffered the same proportional reduction in discriminability as distantly represented images. Distance was determined using a Euclidean approach, measured in activity space; coordinates within this activity space were defined by the activity of multiunits within each pseudo-population. We used the pdist.m function in Matlab to obtain the distance between each pair of images (in z-score-normalized activity space), using control (warm) data. We then cycled through each image and rank-ordered all the other images by their distance to it: the images with the smallest Euclidean distance were ranked first, those with the longest distance last. We then used this ranking to sort the cooling classification accuracy values for that same pair of images. Note that the accuracy values were computed using different data from the data used to compute the distance ranking values. After cycling through all images, this resulted in a matrix of decoding accuracy values, where each row corresponded to an image, and each column corresponded to the accuracy value for its near to far neighbors. We then averaged this matrix to obtain the accuracy-distance curve. We repeated the same process using the simulation data: instead of using the cooling decoding accuracy values, we used the simulated cooling values.

Relevant data and code is available from the authors upon discussion.

## ACKNOWLEDGEMENTS

We thank Gabriel Kreiman for his helpful comments in an earlier draft. This work was funded by NEI grant EY016187 (to MSL) and the Burroughs Wellcome Postdoctoral Enrichment Award (to CRP). Part of this work was realized with assistance from the Core Grant for Vision Research EY12196. Portions of this research were conducted on the Orchestra High Performance Compute Cluster at Harvard Medical School. This NIH-supported shared facility consists of thousands of processing cores and terabytes of associated storage and is partially provided through grant NCRR 1S10RR028832-01. See http://rc.hms.harvard.edu for more information. Part of this work was also conducted with support from Harvard Catalyst | The Harvard Clinical and Translational Science Center (National Center for Research Resources and the National Center for Advancing Translational Sciences, National Institutes of Health Award UL1 TR001102) and financial contributions from Harvard University and its affiliated academic healthcare centers. The content is solely the responsibility of the authors and does not necessarily represent the official views of Harvard Catalyst, Harvard University and its affiliated academic healthcare centers, or the National Institutes of Health.

## AUTHOR CONTRIBUTIONS

CRP and MSL designed the experiments. CRP, SGL and MSL performed the surgical implantations and intraoperative cooling experiments. CRP conducted all following experiments and analyzed the data. CRP wrote the initial manuscript which was then edited by MSL and also approved by SGL.

